# Social dominance and reproduction result in increased integration of oxidative state in males of an African cichlid fish

**DOI:** 10.1101/2022.02.22.481488

**Authors:** Robert J. Fialkowski, Shana E. Border, Isobel Bolitho, Peter D. Dijkstra

## Abstract

Oxidative stress is a potential cost of social dominance and reproduction, which could mediate life history trade-offs between current and future reproductive fitness. However, the evidence for an oxidative cost of social dominance and reproduction is mixed, in part because organisms have efficient protective mechanisms that can counteract oxidative insults. Further, previous studies have shown that different aspects of oxidative balance, including oxidative damage and antioxidant function, varies dramatically between tissue types, yet few studies have investigated oxidative cost in terms of interconnectedness and coordination within the system. Here, we tested whether dominant and subordinate males of the cichlid *Astatotilapa burtoni* differ in integration of different components of oxidative stress. We assessed 7 markers of oxidative stress, which included both oxidative damage and antioxidant function in various tissue types (total of 14 measurements). Across all oxidative stress measurements, we found more co-regulated clusters in dominant males, suggesting that components of oxidative state are more functionally integrated in dominant males than they are in subordinate males. We discuss how a high degree of functional integration reflects increased robustness or efficiency of the system (e.g. increased effectiveness of antioxidant machinery in reducing oxidative damage), but we also highlight potential costs (e.g. activation of cytoprotective mechanisms may have unwanted pleiotropic effects). Overall, our results suggest that quantifying the extent of functional integration across different components of oxidative stress could reveal insights into the oxidative cost of important life history events.

## 1. Introduction

Oxidative cost has been proposed as a potential mediator of life history trade-offs, such as the negative correlation between reproduction and longevity (Alonso-Alvarez et al., 2017; Speakman and Garratt, 2014). Reproduction involves metabolically demanding activities such as egg production, territorial defense, and parental care, leading to increased levels of reactive oxygen species (ROS, also referred to as oxidants). The resulting increased production of ROS, if not sufficiently neutralized by the antioxidant machinery, may lead to increased oxidative stress (Balaban et al., 2005). Given that accumulation of oxidative stress contributes to compromised cell health and disease progression, it is plausible that oxidative stress acts as a constraint in life history decisions, mediating trade-offs between for example fecundity, growth and survival (Dowling and Simmons, 2009; Monaghan et al., 2009; Speakman et al., 2015). Accordingly, studies have shown that reproductive effort can increase oxidative damage and decrease antioxidant capacity in a range of animal species (Christe et al., 2012; Sawecki et al., 2019; Sharick et al., 2015; Stier et al., 2012), and that oxidative stress can negatively affect survival or lifespan (Archer et al., 2013; Bize et al., 2008). In many animals species, social dominance is associated with breeding, and similarly, there is cumulative evidence that social dominance results in an oxidative cost, especially when high dominance is linked to intense agonistic interactions (Beaulieu et al., 2014; Border et al., 2019; van de Crommenacker et al., 2011) and/or increased reproductive effort (Cram et al., 2015; Noguera, 2019; Silva et al., 2018). At the same time, there are also several studies suggesting there is no oxidative cost of social dominance and/or reproduction and that breeding may even reduce oxidative stress (Blount et al., 2016; Costantini et al., 2014; Garratt et al., 2013). This is, in part, due to the fact that organisms have sophisticated defensive mechanisms, such as antioxidant and repair systems, that can avoid or mitigate the negative consequences of oxidative insults (Hõrak and Cohen, 2010; Pamplona and Costantini, 2011). However, few studies have explored the efficiency or robustness of these defensive mechanisms that maintain redox homeostasis.

Maintaining redox balance requires complex integration of various redox components both within and across interconnected cells, organs, and tissues. For example, oxidative insults may activate the transcription factor nrf1 which regulates many detoxifying enzymes and antioxidant genes (Enomoto et al., 2001). Another example of the highly integrative nature of the antioxidant response is the fact that exogenous antioxidant supplementation often leads to compensatory responses such as reduced production of endogenous antioxidants to maintain oxidative balance (Selman et al., 2006). Therefore, assessing the functional integration and interconnectedness of multiple oxidative stress variables could provide important information about the effectiveness of cytoprotective mechanisms that counteract oxidative damage - such as antioxidant responses - as well as the degree of active regulation of redox balance under stressful conditions. For example, hybrids of the newt species *Triturus* exhibited a lower level of functional integration of the antioxidant system compared to the parental species, which was viewed as a potential oxidative cost of interspecific hybridization (Prokić et al., 2018). Likewise, short-term flight in zebra finches led to reduced integration across blood redox markers, which constitutes a potential oxidative cost (Costantini et al., 2013). However, studies examining the effect of challenging life history events on the degree of functional integration of oxidative stress components across different tissue types are largely lacking, with previous studies focusing on only blood or whole body samples (Costantini et al., 2011; Costantini et al., 2013; Prokić et al., 2018).

The cichlid fish *A. burtoni* lives in a lek-like social system in which dominant males defend a spawning territory while subordinate males are non-territorial and are reproductively suppressed (Fernald, 2017). This social structure is reproducible in a laboratory setting by housing several males and females in an aquarium with flowerpot shards acting as spawning territories to encourage territoriality in several males. Dominance hierarchies in *A. burtoni* are characterized by intense rates of aggression with dominant males engaging in continual rank maintenance using border displays and chases (Piefke et al., 2021). Additionally, dominant males show increased reproductive effort as indicated by larger gonads, brighter coloration, increased courtship displays, and higher androgen levels compared to subordinate males (Alward et al., 2020; Border et al., 2018; O’Connell and Hofmann, 2012). We previously found that in *A. burtoni* dominant males had higher levels of plasma oxidative damage (measured as circulating reactive oxygen metabolites) compared to subordinate males (Border et al., 2019; Border et al., 2021; Fialkowski et al., 2021). In the current study, we tested the effect of social status on oxidative stress levels as well as how the integration between different measurements of oxidative stress varies between social states. Markers of oxidative stress included oxidative DNA damage, NADPH-oxidase (NOX) activity (which influences ROS generation), total antioxidant defense and superoxidative dismutase (SOD) activity (more details about these markers can be found in the method section). Most of these markers were measured in different tissue types including blood, liver, muscle, and gonads. We predict that dominant males will have higher levels of oxidative stress (by way of elevated oxidative damage and/or lower total antioxidant capacity) than subordinate males in most tissue types, but we expect this pattern to be redox marker- and tissue-specific. We also predict that dominant and subordinate males differ in the degree of integration between different measurements of oxidative stress.

## 2. Methods

### 2.1 Animals and housing

The cichlid *Astatotilapia burtoni* used in this study were descended from a wild-caught stock population from Lake Tanganyika. Fish were housed in aquaria kept at 28°C with a 12-h light/dark cycle and 10 min each dusk and dawn period to mimic natural settings. Aquaria contained gravel substrate and terracotta shelters. Fish were fed a combination of cichlid flakes (Omega Sea LLC) and granular food (Allied Aqua) each morning. Continuous water flow and central mechanical and biological filtration occurred throughout the entirety of the experiment. All fish used in this experiment were initially housed as larvae and juveniles in 110-L tanks until they were approximately 4 months of age, after which they were transferred to a 407-L tank until randomly selected individuals were transferred to experimental tanks. All fish used in this experiment were adults and had been raised in mixed-sex groups. Experimental males were tagged just below the dorsal fin with colored beads attached to a plastic tag using a stainless-steel tagging gun (Avery-Dennison, Pasadena, CA). All procedures were approved by Central Michigan University Institutional Animal Care and Use Committee (IACUC protocol 15-22).

### 2.2 Experimental design

We divided experimental 110-L tanks in half widthwise with clear, perforated acrylic barriers to create two compartments as described elsewhere (Fialkowski et al., 2021). In each experimental compartment we placed a group comprised of three males and five females (*n*=31 groups). Fish were between 10 – 12 months of age. Males were weighed and their standard length (SL) was measured before adding them to the experimental tanks. We provided one flowerpot shard in each compartment which was occupied by the dominant male in each group. In most groups (27 out of 31) the largest male (at least 0.1 g bigger than the other males) attained social dominance while the other two males in each group became subordinate. Hence, social status was mostly, albeit not perfectly, assigned based on size asymmetry. The initial body mass of dominant males was 5.2 ± 0.2 g (range 2.6 – 6.8, *n*=31) and subordinate males was 4.2 ± 0.1 g (range 3.0 – 6.0, *n*=31). Given the considerable size overlap between social states *across* groups, size is unlikely a confounding factor. Fish were able to interact physically with members in their own group and visually with those in the adjacent compartment, allowing for the full expression of dominant behaviors, including aggressive interactions between dominant males between compartments. We collected tissue and blood from the dominant male and one randomly selected subordinate male after housing fish in this arrangement for 6 – 7 weeks.

### 2.3 Behavioral observations

We recorded male social status (dominant or subordinate) three times per week following a two week stabilization period as described previously (Border et al., 2019). In brief, dominant males were brightly colored, defended a flowerpot and had a dark eye bar while subordinate males are cryptically colored and shoal with females. We filmed each group in the morning before 10 a.m. weekly during the final 4 weeks of the experiment, with the final recording made on the morning of tissue collection. For each group, we quantified the behaviors of the focal males (dominant male and subordinate male for which we collected tissue) over a five-minute period. One person scored the frequency of chases, lateral displays, border displays, cave visits, courtship displays, and flees as previously described (Fialkowski et al., 2021). We were unable to carry out the observations blind with respect to social status due to the readily apparent differences in coloration and behaviors observed between dominant and subordinate males. Behavioral coding was completed using Behavioral Observation Research Interactive Software (BORIS)(Friard and Gamba, 2016).

### 2.4 Tissue sampling methods

Immediately following the final behavioral observation, we removed focal males (one dominant and one subordinate male) to collect blood and tissue. Males were weighed and their standard length was measured before blood was drawn. Blood time for each male was measured as the time from initial disruption of the tank (when the lid was removed) to the completion of blood collection for that individual ranging from 2.5 −11.75 minutes, with individuals processed in a randomized order with respect to social status. We collected approximately 25-100 μL of blood from each male (*n*=31 each). Blood was drawn through the dorsal aorta using heparinized 26-gauge butterfly needles (Terumo) and transferred to heparinized centrifuge tubes that were placed on ice.

Immediately after blood was drawn, males were euthanized via rapid cervical transection and tissues (gonads, liver, muscle) were collected and flash frozen in liquid nitrogen. Gonads were weighed prior to freezing. Frozen tissue samples were immediately placed in 2 mL tubes on a M15 Coolrack block (Biocision, Larkspur, CA, USA) on dry ice. Tissue collection time (measured as the time from initial disruption of the tank to freezing tissue) ranged from 3.1 – 23.0 minutes. Blood samples were centrifuged for 10 minutes at 4000g before plasma red blood cells were separated. Blood cells, plasma and tissue samples were stored at −80°C until used for analysis.

### 2.5 Measurement of oxidative stress

#### 2.5.1 Choice of markers of oxidative stress

As an overall marker of oxidative stress, we selected plasma reactive oxygen metabolites (ROMs). ROMs are primarily organic hydroperoxides which include a range of oxidized substrates, such as polyunsaturated fatty acids, proteins and nucleic acids, and thus represents a comprehensive measure of oxidative damage (Costantini, 2016). We measured plasma antioxidant capacity in three ways: total antioxidant capacity (TAC), the OXY-adsorbent Test (OXY), and the Biological Antioxidant Potential Test (BAP). TAC is a cumulative measure of total antioxidants based on the ability of antioxidants with both low molecular weight and high molecular weight (such as enzymatic antioxidants) to scavenge peroxyl radicals using the oxygen radical absorbance capacity (ORAC) assay (Marrocco et al., 2017)). Like TAC, OXY also measures total antioxidant capacity but it is based on a different biochemical reaction (it measures the ability of plasma to withstand oxidative insult from hypochlorous acid) and tends not to measure larger, enzymatic antioxidants. The OXY assay was selected because it is a widely used marker for systemic levels of total antioxidant capacity including in haplochromine cichlid fish (Dijkstra et al., 2011; Dijkstra et al., 2016). The BAP test is based on iron oxidation and provides insight into more dynamic antioxidant activity of substances with lower molecular weight. In addition to plasma TAC, we measured TAC in liver, gonad, and muscle using the ORAC assay. We also measured superoxide dismutase (SOD) activity, an important enzymatic antioxidant that removes oxygen radicals via conversion to molecular oxygen and uric acid. SOD was measured it in the liver and gonads. Oxidative DNA damage was evaluated by measuring levels of 8-hydroxy-2’-deoxyguanosine (8-OhDG) in red blood cells, liver, and gonads. As an indirect marker of ROS production, we measured nicotinamide adenine dinucleotide phosphate-oxidase (NADPH-oxidase or NOX) activity in the liver and gonads. NOX are a major source of superoxide (a primary reactive oxygen species) in various tissue cell types (Bedard and Krause, 2007). We were unable to measure muscle NOX and muscle SOD activity due to technical difficulties (NOX) or lack of sample (SOD). We measured several redox markers in gonadal tissue since we expected social status differences in the reproductive system. The liver is particularly vulnerable to oxidative damage and is frequently included in tissue oxidative stress analyses. Muscle was also selected because of its importance for physical activities.

Here we consider increases in (potential) oxidative damage and/or lower antioxidant capacity as indicative of increased oxidative stress. More specifically, increased DNA damage and NOX activity may reflect elevated oxidative stress while lower total antioxidant protection (TAC, OXY, and BAP) could reflect antioxidant depletion under high oxidative stress levels. However, it is important to note that increased ROS production can lead to compensatory mechanisms, such as upregulated SOD. It is therefore difficult to interpret SOD findings in relation to the level of oxidative stress an organism experiences (Monaghan et al., 2009).

All samples were run in duplicate and included a pooled sample to evaluate interplate variability with the exception of SOD. For all assays we used clear flat-bottom 96-well plates unless indicated otherwise. For all assays, the intra-assay CV (coefficient of variation) and the inter-assay CV were typically below 5% and 13%, respectively.

#### 2.5.2 Protein quantification

The protein concentration of each prepared tissue sample (tissue TAC, SOD, and NOX activity) was measured with a Bicinchoninic Acid (BCA) Protein Assay kit (Pierce, Rockford IL) following the manufacturer’s protocol. Frozen supernatant was thawed on ice and diluted 1:4 with buffer used in their respective sample preparation. We used 10 μL of this diluted supernatant in the assay. Absorbance was read by a plate reader (Epoch2T, Biotech Instruments, Winooski, VT, USA).

#### 2.5.3 Circulating Reactive Oxygen Metabolites (ROMs)

We measured the concentration of plasma ROMs (primarily organic hydroperoxides) in blood plasma using the widely used d-ROM test (Diacron, Grosseto, Italy) using 4 μL plasma per well as previously described (Border et al., 2019). We calculated values in Carratelli units which was then converted to H_2_O_2_ mg/dL. Absorbance was read by a plate reader (Epoch2T, Biotech Instruments, Winooski, VT, USA).

#### 2.5.4 Circulating Antioxidants

##### Total Antioxidant Capacity (TAC)

Circulating TAC was determined in diluted blood plasma (1:100) in 7.4 pH PBS using 20 μL of diluted sample via an oxygen radical absorbance capacity (ORAC) assay as previously described (Border et al., 2019). For circulating TAC, samples were reported as μmol TE/dL of the sample. Absorbance was read by a plate reader (Spectramax M3, Molecular Devices, Sunnyvale, CA, USA).

##### OXY-adsorbent Test (OXY)

For the OXY assay (Diacron, Grosseto, Italy), we diluted blood plasma (1:100) with distilled water using 2 μL of diluted sample per well following the protocol as described previously (Dijkstra et al., 2016). Plasma OXY concentration was expressed as μmol of HClO/mL of sample. Absorbance was read at 550 nm by a plate reader (Spectramax M3, Molecular Devices, Sunnyvale, CA, USA).

##### Biological Antioxidant Potential Test (BAP)

For the BAP assay (Diacron, Grosseto, Italy), we used 2 μL of plasma for each well and followed the manufacturer instructions. Plasma BAP concentration was expressed as μmol/mL of sample. Absorbance was read at 505 nm by a plate reader (Spectramax M3, Molecular Devices, Sunnyvale, CA, USA).

#### 2.5.5 Tissue Total Antioxidant Capacity (TAC)

TAC was determined in tissue (liver, gonad, and muscle) following the same ORAC procedure as described for plasma with the following exceptions for all tissues. Tissues were removed from −80°C and homogenized on ice in 0.250 mL lysis buffer (20 mM Tris–HCl, 137 mM NaCl, 1% NP-40, 10% glycerol, 2 mM EDTA) using an Omni Tissue Master (Omni International, Kenosha, Wisconsin), then centrifuged at 4°C at 17,000g for 10 minutes. Supernatant was collected and used to run BCA and ORAC. For ORAC, protein concentrations were standardized to ~150 μg/mL and reported as μmol TE/μg protein. Absorbance was read by a plate reader (Spectramax M3, Molecular Devices, Sunnyvale, CA, USA).

#### 2.5.6 Superoxide dismutase (SOD)

SOD was measured using Water Soluble Tetrazolium Salts (WSTs) via a modified competitive assay (Peskin and Winterbourn 2000) in liver and gonads. Samples were taken from −80°C and homogenized on ice in a PBS (pH 7.4, 75mM). Tissue homogenates were then centrifuged at 4 °C at 10,000 g for 10 min. The supernatant was then transferred to a new tube and stored at −80 °C until used for analysis. Protein concentration was measured using BCA before preparing each supernatant at multiple protein concentrations (liver: 0.375 μg/μL, 0.25 μg/μL, 0.125 μg/μL, and 0.05 μg/μL; gonads: 0.4 μg/μL, 0.3 μg/μL, 0.2 μg/μL, and 0.1 μg/μL). SOD activity was measured as described previously (Fialkowski et al., 2021). The amount of SOD content was reported per μg of protein (one unit of SOD is defined as the amount of enzyme in 20 μL of the sample solution that inhibits the reduction reaction of WST-1 with superoxide anion by 50%). Absorbance was read by a plate reader (Epoch2T, Biotech Instruments, Winooski, VT, USA).

#### 2.5.7 NADPH-Oxidase (NOX) activity

NOX activity was measured in the liver and the gonads via a lucigenin-based chemiluminescence assay using a microplate luminometer (Fialkowski et al., 2021). Liver samples were taken from −80°C and homogenized on ice in a Krebs-HEPES buffer. The homogenized samples (before centrifugation) were subjected to a freeze-thaw cycle to ensure total cell lysis. After thawing, samples were centrifuged at 4°C at 600g for 10 minutes and supernatant was transferred and stored at −80°C. For gonad samples, the supernatant was subjected to an additional freeze-thaw cycle to ensure total cell lysis. For each sample, an aliquot of supernatant was used to run BCA. To measure NOX activity, samples containing 30 μg (liver) or 15 μg (gonad) of protein were placed in a solid black 96-well plate (Corning 3912), then the reaction was initiated out of direct light by the addition of buffer containing lucigenin and NADPH, with a final concentration of 5uM lucigenin and 100uM NADPH. The plate was kept covered for transport to luminometer (Tecan infinite F200 Pro, Tecan Life Sciences, Männedorf, Zürich, Switzerland) and readings of Relative Light Units (RLU) were taken every 2 minutes for 20 minutes. The area under the curve was calculated for minutes 2-20, and the results were expressed as RLU per minute per μg protein after subtraction of background chemiluminescence.

#### 2.5.8 Oxidative DNA damage

Oxidative DNA damage was evaluated for 8-OhDG damage using a DNA damage ELISA kit (StressMarq Biosciences Inc.) (Fialkowski et al., 2021). DNA was extracted from packed red blood cells (PRBCs) and frozen tissue samples (liver, gonad) using a commercially available DNA extraction kit (Zymo quick-DNA miniprep plus kit) as described previously. Extracted samples were stored at 4 °C until digestion. Samples were digested at a standardized concentration of ~200 ng/ul (PRBCs, gonad) or 400-500 ng/ul (liver) using a modified digest mix protocol by Quinlivan and Gregory (2008) and stored at −20 °C until use in DNA damage plate (Quinlivan and Gregory, 2008). Digested samples were tested for 8-OhDG damage using a DNA damage ELISA kit at 12x dilution for blood and gonad samples and a 10x dilution for liver samples, with all samples run in duplicate following the manufacturer’s instructions. Absorbance was measured using a microplate reader (Epoch2T, Biotech Instruments, Winooski, VT, USA). 8-OhDG concentration was standardized relative to total DNA concentration and reported in ng/μL.

#### 2.5.9 Testosterone levels

To confirm that dominant males upregulated their reproductive system, we quantified circulating testosterone levels using competitive ELISA kits (Enzo Life Sciences) as previously described (Border et al., 2019). Absorbance was read by a plate reader (Epoch2T, Biotech Instruments, Winooski, VT, USA).

### 2.6 Statistical analysis

We calculated a dominance index score for each weekly 5-minute focal observation as the sum of (aggressive behavior + reproductive behaviors) - fleeing events per min, as done previously in *A. burtoni* (Maruska et al., 2013). Gonadosomatic index (GSI) was calculated as (gonad weight/total body weight)*100. Specific growth rate was expressed as the daily percentage weight change from the beginning to the end of the experiment relative to the initial weight (calculated as [ln(body weight_final_) - ln(body weight_initial_)] x 100/days) (Ricker, 1975). Since males that were initially smaller grew more during the experiment, we calculated the residuals of specific growth rate using a linear regression and used this as our growth variable in the analysis.

All analyses were conducted in R v3.4.3. We analyzed our data using the R packages lme4, lmerTest, MASS (Bates et al., 2015), and glmmTMB (Brooks et al., 2017). We identified and excluded outliers based on Tukey’s rule (between 0 and 2 values were excluded per measurement). In addition, samples size for oxidative stress measurements varied depending on availability of tissue or technical constraints (Table 1). We used linear mixed models (LMMs) with a maximum-likelihood protocol. For count and proportional data, we used generalized linear mixed models (GLMMs). In each model, we used ‘pair code’ as random effect to account for fish that were housed in the same group. To evaluate the validity of our LMM and GLMM models, we examined the residuals, qqplots, and plots of predicted values versus residuals. We report mean ± SE for our model estimates.

**Table 1.** Statistical results comparing measurements of oxidative stress between dominant and subordinate males. Significant effects are shown in bold.

| 710 **Marker** | **Estimate** | **df** | **t value** | **P value** | **Sample sizes** |  |
| --- | --- | --- | --- | --- | --- | --- |
|  |  |  |  |  | **Dominant** | **Subordinate** |
| 711 Plasma ROMs | -0.603±0.191 | 26.46 | -3.167 | **0.00385\*\*** | 29 | 29 |
| 712 Plasma TAC | -95.52±62.03 | 58.00 | -1.54 | 0.129 | 28 | 30 |
| 713 Plasma OXY | -35.24±42.13 | 38.00 | -0.836 | 0.408 | 19 | 19 |
| 714 Plasma BAP | -1812.9±1083.6 | 26.0 | -1.673 | 0.106 | 13 | 13 |
| 715 Liver TAC | 0.005±0.068 | 31.00 | 0.07 | 0.944 | 31 | 31 |
| 716 Gonad TAC | -0.058±0.051 | 30.28 | -1.151 | 0.259 | 31 | 30 |
| 717 Muscle TAC | 0.069±0.068 | 61.00 | 1.021 | 0.311 | 30 | 31 |
| 718 Blood DNA damage | 0.011±0.011 | 0.25 | 1.008 | 0.323 | 29 | 28 |
| 719 Liver DNA damage | -0.0034±0.0060 | 0.18 | -0.565 | 0.579 | 26 | 19 |
| 720 Gonad DNA damage | -0.0135±0.011 | 0.27 | -1.246 | 0.223 | 29 | 26 |
| 721 Liver NOX | -4.36±2.48 | 28.88 | -1.755 | 0.0899 | 29 | 29 |
| 722 Gonad NOX | -6.68±3.05 | 30.96 | -2.19 | **0.0362\*** | 30 | 30 |
| 723 Liver SOD | -0.011±0.036 | 49.00 | -0.313 | 0.755 | 26 | 23 |
| 724 Gonad SOD | -0.023±0.029 | 27.40 | -0.79 | 0.436 | 27 | 20 |

#### 2.6.2 Analysis of behavior, GSI, and testosterone

To confirm the assigned social status of each male, we compared behavior, GSI, and testosterone levels between dominant and subordinate males. We tested for differences between social states in dominance index using a LMM. Shoaling duration was analyzed using a GLMM assuming a Gaussian distribution with log link function and foraging bouts were analyzed using GLMM assuming a negative binomial distribution. In addition to ‘pair code’, we used ‘fish code’ nested within ‘pair code’ as random effect to account for the 4 weekly measurements of behavior for each focal male. GSI was analyzed using a LMM and testosterone levels were analyzed using a GLMM assuming a Gaussian distribution with log link function.

#### 2.6.3 Analysis of oxidative stress

Measurements of oxidative stress were compared between dominant and subordinate males using LMMs. To examine whether covariance patterns across the different oxidative stress measurements varied by social status, we created clustered correlation matrices for dominant and subordinate males separately. We then carried out a hierarchical cluster analysis to identify clusters of oxidative stress measurements that were coregulated. We obtained *P*-values using multiscale bootstrap resampling from the pvclust package (Suzuki and Shimodaira, 2006). Since growth rate and investment in territorial defense and reproduction could influence oxidative state, we also carried out the same hierarchical cluster analysis after adding GSI, specific growth rate, dominance index (based on the final five-minute focal observation, closest to tissue sampling) and testosterone to the oxidative stress dataset.

## 3. Results

### 3.1 Males become dominant or subordinate

Males did not change social status a during the entire duration of the experiment (Fig. 1A). The dominance index (the difference between aggressive behavior and fleeing events) was significantly higher in dominant males (LMM, −12.6871±0.4730, t_62_=-26.82, *P*<0.00001). Subordinate males spent more time shoaling than dominant males (GLMM, 2.59±0.50, z=5.19, *P*<0.00001). Social dominance was linked to more reproductive behavior, with dominant males showing more courtship behavior than subordinate males (GLMM, zero-inflation model, 3.27±0.61, z=5.34, *P*<0.00001). We observed mouthbrooding females in all groups during the entire duration of the experiment, suggesting that all dominant males spawned (females typically spawn with dominant males, PDD pers. obs.).

**Fig. 1.**
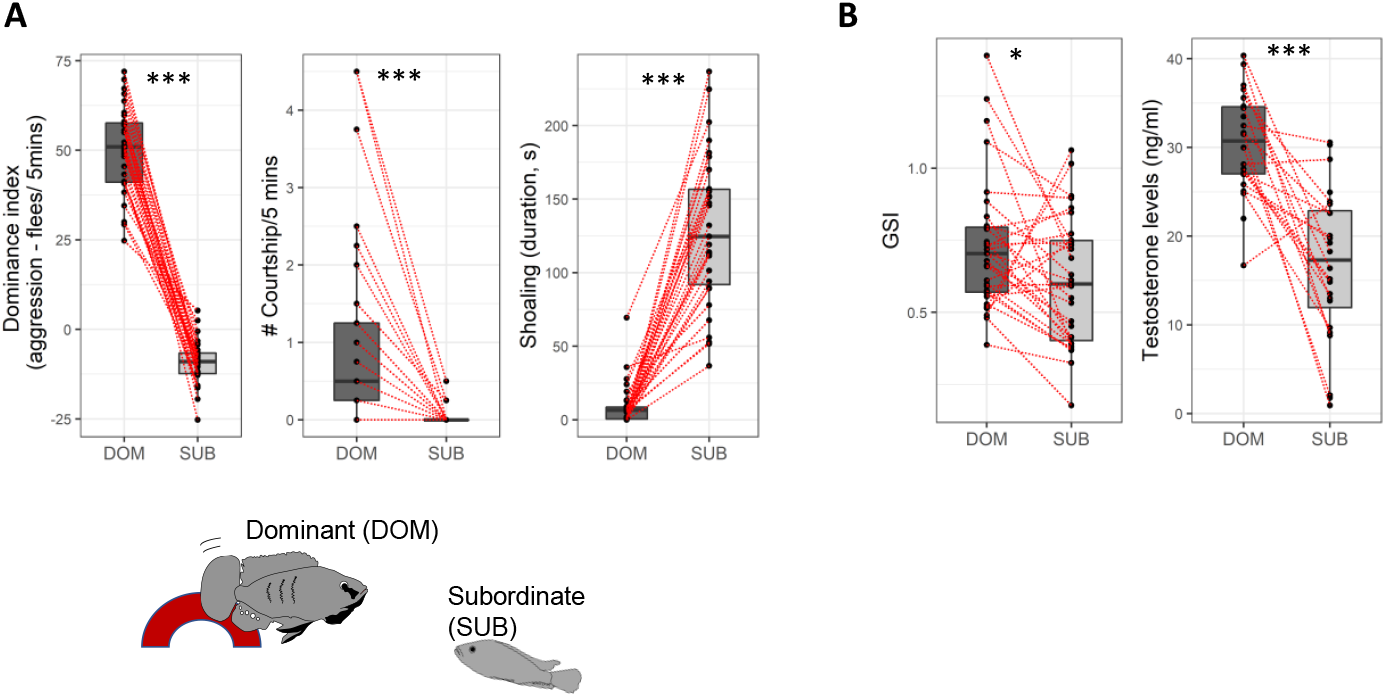
Levels of behavior, testosterone, and gonadosomatic index by social status. (A) Dominant males show more aggressive and courtship behaviors while subordinate males show more subordinate behaviors such as fleeing and shoaling during four weekly observations. Shown are the rates (dominant index per minute, courtship per 5 minutes, shoaling amount of time spent) averaged across the final four weeks prior to tissue sampling. (B) Dominant males also had higher gonadal somatic index (GSI) and higher circulating testosterone levels than subordinate males. Bold lines indicate medians. Boxes enclose 25th to 75th percentiles. Error bars enclose data range, excluding outliers. Dots are data points and red lines connect data for males that were housed together. * *P*<0.05, *** *P*<0.001

Dominant males had higher gonadosomatic index (LMM, −0.125±0.050, t_31_=-2, *P*=0.0189) and circulating testosterone levels (GLMM, −0.595±0.102, z=-5.85, P<0.00001) than subordinate males, confirming that the former had an activated reproductive system (Fig. 1B). Finally, dominant males grew faster than subordinate males (specific growth rate, dominant males: 1.044±0.067, subordinate males: 1.019±0.063) and after correcting for the effect of initial body weight this effect of social status on growth rate was significant (LMM, −0.196±0.067, t_31_=-2.94, *P*=0.006).

### 3.2 Dominant males experienced greater circulating oxidative damage than subordinate males

We compared a total of 14 oxidative stress measures between dominant and subordinate males (Fig. 2, Table 1). Plasma reactive oxygen metabolites (ROMs), a marker of overall oxidative damage, was higher in dominant males than in subordinate males (Table 1), consistent with previous findings in the same species (Border et al., 2019; Fialkowski et al., 2021). Dominant males displayed higher NOX activity in the gonads and liver, although this effect was only significant in the gonads (Table 1). There were no significant status differences in the other measurements of oxidative stress (Table 1).

**Fig. 2.**
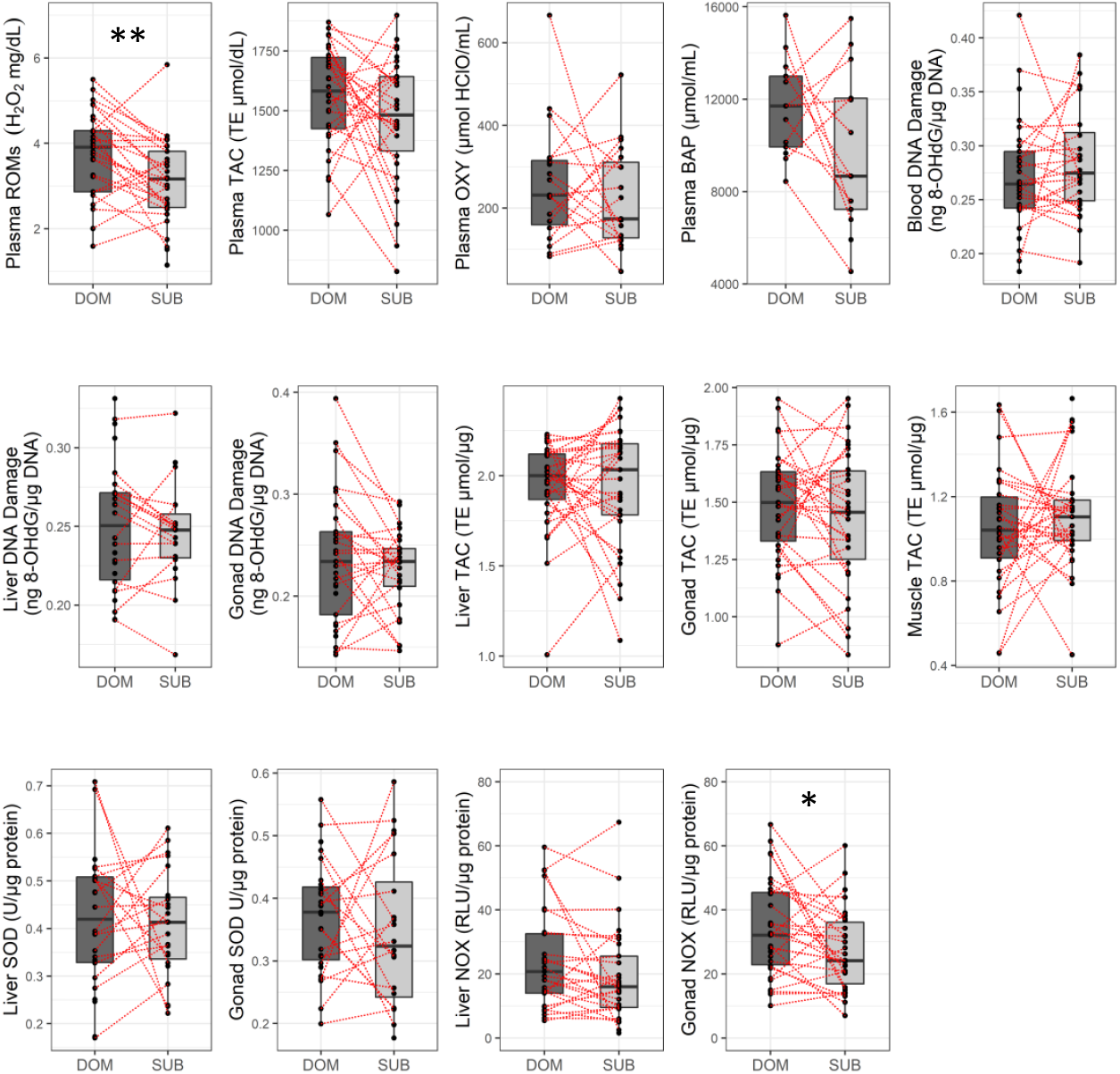
Measurements of oxidative stress by social status. Reactive oxygen metabolites (ROMs) and NOX (NADPH-oxidase) activity were higher in dominant males than in subordinate males. The other markers did not vary by social status. Boxes enclose 25th to 75th percentiles. Error bars enclose data range, excluding outliers. Dots are data points and red lines connect data for males that were housed together. For statistics, see table 1. * *P*<0.05, ** *P*<0.01

### 3.3 Dominant males express greater covariance patterns of oxidative stress profile than subordinate males

We tested whether dominant and subordinate males vary in co-variance patterns across the different markers of oxidative stress and tissue types by examining clustered correlation matrices for dominant and subordinate males separately (Fig. 3A). In dominant males, significant clusters included both antioxidant function and oxidative damage across multiple tissue types while in subordinate males only one plasma antioxidant cluster was significant. Specifically, there were three significant clusters in dominant males, one involving plasma BAP, liver TAC, and gonad SOD, a second cluster involving muscle TAC and liver SOD, and a third cluster comprised of the remaining variables (all *P*<0.05). By contrast, there was only one small cluster that was significant in subordinate males comprised of plasma BAP and plasma OXY (*P*<0.05).

**Fig. 3.**
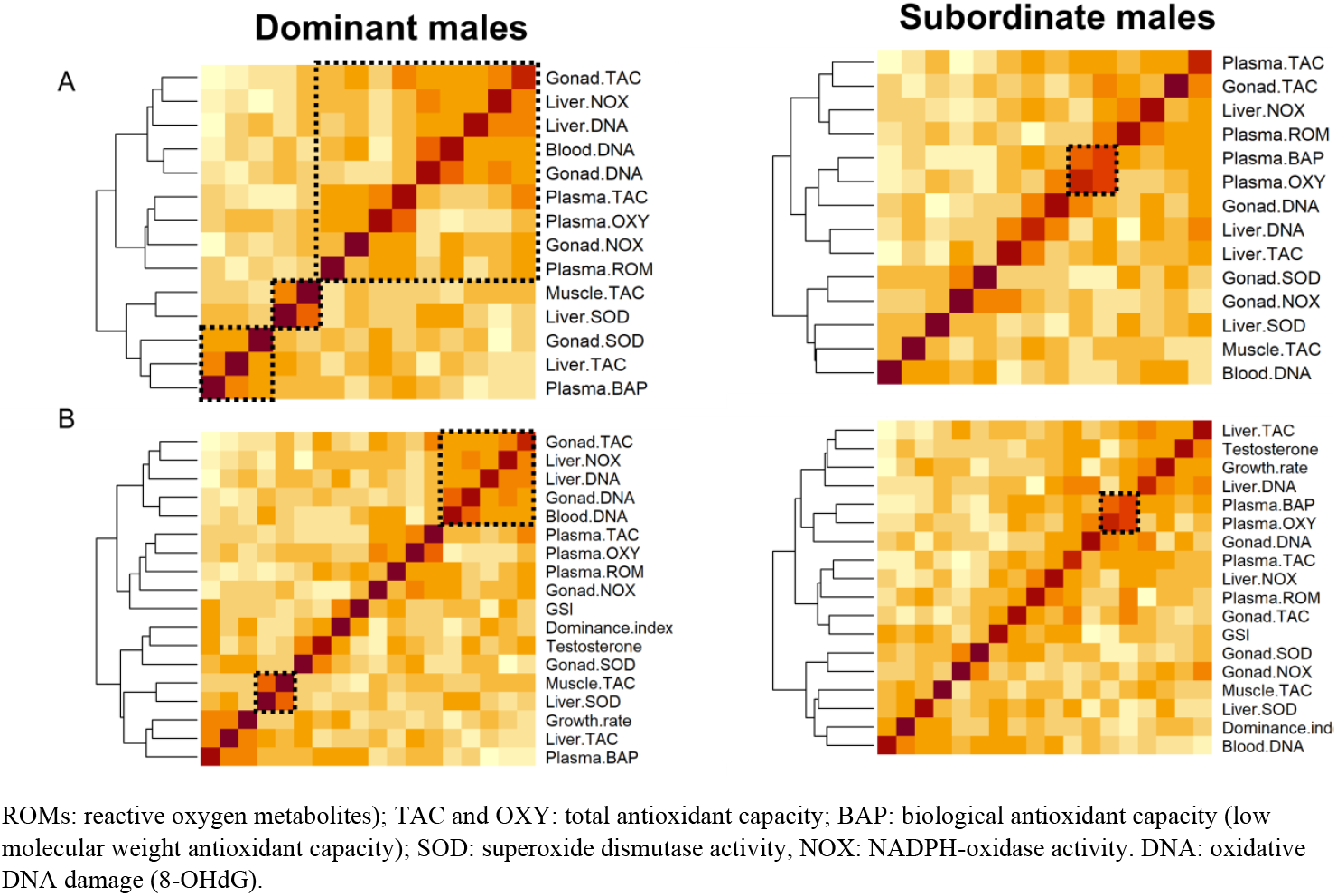
Covariances patterns across markers of oxidative stress by status. (A) Covariance across 14 different oxidative stress measurements for dominant and subordinate males. (B) Covariance patterns for the same oxidative stress measurements combined with indicators of reproduction and social dominance (dominance index, gonadosomatic index (GSI), and testosterone). Hierarchical clustering revealed more significant clusters (indicated by dashed box) in dominant males than in subordinate males. Each cell represents the correlation value, with hotter colors representing a more positive correlation.

Given that investment in growth, social dominance, and reproduction may be linked to oxidative stress, we also tested for co-variance patterns in oxidative stress datasets that included GSI, specific growth rate, testosterone, and dominance index. This analysis revealed two significant clusters in dominant males versus only one in subordinate males (Fig. 3, all *P*<0.05). Similar to the analysis that only included measurements of oxidative stress, there were more clusters in dominant males than subordinate males. Clusters in dominant males included both antioxidant function and oxidative damage across tissue types, supporting that social dominance is causally linked to a higher degree of functional integration of oxidative state.

## 4. Discussion

We found clear dominance hierarchies with dominant males having upregulated GSI and higher testosterone levels than subordinate males. However, we found limited evidence that dominant males had higher levels of oxidative stress than subordinate males when comparing independent oxidative stress measurements. Importantly, there were more significant clusters of coregulated oxidative stress variables in dominant males than in subordinate males, suggesting that dominant males have more efficient or more active regulation of oxidative balance than subordinate males. Below, we discuss these findings in more detail.

### 4.1 Limited cost of social dominance when markers are evaluated in isolation

Our findings suggest that social status-specific differences in oxidative balance is highly tissue- and marker-specific. Organisms are comprised of a complex set of integrated organs performing unique functions. As a result, energetically demanding activities such as reproduction and defending high rank likely affect parts of the body differently (Costantini, 2019; Speakman and Garratt, 2014). It is therefore unsurprising that reproduction may elevate, reduce, or have no impact on oxidative damage and/or antioxidant function depending on which macromolecules and tissues are considered, which our findings here support (Garratt et al., 2011; Garratt et al., 2013; Ołdakowski et al., 2015; Yang et al., 2013). However, it was surprising that when comparing markers of oxidative stress in isolation, dominant and subordinate males only differed in two measurements of oxidative stress. We also note that increased NOX activity in the gonads of dominant males is consistent with higher NOX signaling in mature sperm, which does not necessarily constitute a ‘cost’ given the important role of redox signaling in sperm function (Tremellen, 2012). Furthermore, there was no social status effect on oxidative DNA damage, even though we measured it in three different tissue types.

There are several explanations for the limited status-dependent differences in oxidative stress, including the ability to effectively manage or minimize oxidative stress and the fact that our animals were housed under benign lab conditions. Since oxidative stress is an ever present problem, organisms have evolved multi-faceted highly-regulated cytoprotective mechanisms to mitigate oxidative damage (Balaban et al., 2005). We recently induced social status transition from subordinate to dominant position in *A. burtoni* and found that social ascent was associated with dynamic changes in plasma ROMs, plasma TAC, liver TAC and liver SOD activity (Fialkowski et al., 2021). Specifically, plasma TAC was rapidly depleted while liver SOD and liver TAC were increased during social ascent. However, after 2 weeks of social dominance, dominant males did not have different levels of liver SOD or TAC and had higher plasma ROMs than subordinate males, consistent with the current study.

The fact that in the current study we did not detect social status differences in oxidative balance in most tissue types could be due to a limited cost of social dominance when the hierarchy is stable (for a study comparing stable hierarchies and social ascension in another cichlid species, see (Culbert et al., 2022)). Our findings are consistent with the notion that animals undergo behavioral and physiological adjustments to cope with demanding life history activities. For example, the onset of reproduction is often associated with dramatic metabolic and morphological remodeling to cope with the energetic demands of reproduction (Reiff et al., 2015) and these changes may also involve upregulated antioxidant defense (Blount et al., 2016). Further, our experiments were carried out in captivity under benign conditions providing ad libitum food and protection. It is possible that the cost of social dominance is more pronounced when dominant individuals face additional challenges, such as parental care, food shortage, temperature stress, parasites, or social instability (Speakman et al., 2015). For example, in the white-browed sparrow weaver (*Plocepasser mahali*), highly ranked females but not highly ranked males experienced increased oxidative stress after breeding, presumable due to the fact that females exert more effort during reproduction in relation to egg laying, incubation and egg provisioning (Cram et al., 2015).

### 4.2 Social status differences in integration of oxidative stress components

Social status predicted the extent of functional integration across different components of oxidative stress. Across all 14 measurements of oxidative stress (i.e. all markers and tissue types), we observed three significant co-regulated clusters in dominant males and only one cluster in subordinate males. In dominant males, all oxidative stress variables were part of one of these clusters. In addition, in dominant males all clusters contained variables from different tissue types, in contrast to the situation in subordinate males (in the latter, plasma BAP and plasma OXY formed a cluster, which is not surprising given that both measure overlapping components of antioxidant capacity in blood). Further, in dominant males there was coregulation between different measurements within the same tissue as well as the same measurement across different tissue. It was particularly interesting to note that all oxidative DNA damage measurements were included in this cluster. The other two smaller clusters in dominant males linked different measures of antioxidant function (TAC and SOD) across different tissue types, both containing measures in liver linked to either gonads or muscle. It is difficult to interpret the functional significance of these smaller clusters in dominant males, but they suggest a relatively high level of coordination between different components of the antioxidant defense system across completely different organs.

The more modular redox responses in dominant males relative to subordinate males could be an indication of a more efficient and/or active management of oxidative balance. This finding suggests that dominant males pay a reduced cost to maintaining oxidative balance due to more effective neutralization of oxidative insults by antioxidant defense systems or dominant males benefiting from a more robust, stable system guarding oxidative balance relative to subordinate males. The lack of modularity in subordinate males could suggest that social subordination is associated with increased dysregulation of oxidative balance, and perhaps increased (and costly) investment into maintaining redox balance. This notion that a low level of integration reflects an oxidative cost is consistent with lower integration of antioxidant parameters observed in hybridizing newts as a cost of interspecific hybridization (Prokić et al., 2018) and exercise-induced loss of integration in zebra finches (Costantini et al., 2013). It is also supported by proteomic and metabolomic studies suggesting that the degree of integration or connectivity in metabolic networks may reveal information about the robustness or efficiency of systems that maintain stability and homeostasis. Integration of these systems may decline with age due to failure in communication between interacting units (Hoffman et al., 2017).

However, increased modularity observed in dominant males may also entail costs. More integration may be a manifestation of more active management of oxidative balance. Although the relative energetic cost of upregulating antioxidant enzymes is probably low, active regulation of redox balance is not cost-free due to ‘physiological constraints’ or pleiotropic effects of activating antioxidant response systems (Pamplona and Costantini, 2011). For example, in addition to being a damaging by-product, ROS also have important cell signaling functions, and mounting an antioxidant response could also quench ROS that have beneficial effects (Linnane et al., 2007). Consequently, upregulation of antioxidant enzymes may lead to detrimental side effects (Barajas et al., 2011). The interpretation of increased integration of oxidative stress in dominant males relative to the oxidative cost of social dominance/reproduction is complicated, and future studies should shed more light on functional significance of variation in integration in the context of life history trade-offs. Specifically, to what extent does the degree of integration reflect robustness and efficiency of interacting systems that maintain redox homeostasis? How is the relationship between integration and robustness/efficiency modulated by the type, duration, and magnitude of stressors associated with social dominance and reproduction? And based on this information on the link between integration and efficiency/activation of the system, what are the long-term fitness consequences? These are interesting questions that are not always easy to tackle (e.g. addressing some of these questions requires longitudinal sampling of the same individuals, which is challenging unless non-invasive sampling techniques are used (Alonso-Alvarez et al., 2017)) but may prove useful in future studies.

To conclude, we found that dominant *A. burtoni* males experienced more oxidative stress in only two oxidative stress markers (plasma ROMs and gonad NOX activity) out of a total of 14 different oxidative stress measurements. We found evidence for more integrated redox regulation, and hence more active or efficient management of oxidative balance in dominant males. Whether this supports the oxidative cost of social dominance remains to be tested in future studies. We propose that studies on the oxidative cost of demanding life history events more generally (e.g. migration: (Eikenaar et al., 2020); parental care: (Guindre-Parker and Rubenstein, 2018)) should include analyses of how different components of oxidative stress are interconnected. Even in the absence of difference in mean values in redox markers between for examples breeders and nonbreeders, there might be differences in correlation structure that could provide important insights into the role of oxidative stress as a mediator of life history trade-offs.

## Author’s contributions

Authors PD and SB contributed to the conception and design of the experiment. RF, SB, and PD contributed material preparation and data collection with IB performing behavioral coding. The first draft of the manuscript was written by PD and all authors commented on previous versions of the manuscript. All authors read and approved the final manuscript.

## Ethics approval

All procedures were approved by the Institutional Animal Care and Use Committee (IACUC, protocols #15-22 and #18-10) prior to conducting the experiment. All applicable international, national, and/or institutional guidelines for the use of animals were followed.

## Acknowledgements

We thank Ross DeAngelis and members of the Dijkstra lab for providing helpful comments to earlier drafts of the manuscript. We would also like to thank Hannah Janeski for technical assistance. Deric Learman and Benjamin Swarts are thanked for allowing us to use the plate readers in their lab. Travis Moore assisted us with the behavioral analysis.

